# Increased plasma lipopolysaccharide-binding protein and altered inflammatory mediators in overweight women suggest a state of subclinical endotoxemia

**DOI:** 10.1101/2023.05.18.540879

**Authors:** Christine N. Metz, Xiangying Xue, Prodyot K Chatterjee, Robert P. Adelson, Jesse Roth, Michael Brines, Kevin J. Tracey, Peter K. Gregersen, Valentin A. Pavlov

**Affiliations:** The Feinstein Institutes for Medical Research, Northwell Health, Manhasset, NY 11030, USA; Zucker School of Medicine at Hofstra/Northwell-Hofstra University, Hempstead, NY 11550, USA

**Author notes:** Corresponding authors: Christine Metz, PhD -, Valentin Pavlov, PhD –.

**Keywords:** inflammation, overweight, women, endotoxemia, cytokines, adipokines, cardiovascular disease

## Abstract

Chronic low-grade inflammation has been recognized as an underlying event linking obesity to cardiovascular disease (CVD). However, inflammatory alterations in individuals who are overweight remain understudied. To provide insight, we determined the levels of key circulating biomarkers of endotoxemia and inflammation, including lipopolysaccharide-binding protein (LBP), CRP, IL-6, leptin, and adiponectin in adult female subjects (n=40) who were lean or overweight and had high cholesterol and/or high blood pressure - two important conventional risk factors for CVD. Plasma levels of LBP were significantly higher in the overweight group compared with the lean group (P=0.005). The levels of CRP were also significantly higher in overweight subjects (P=0.01), as were IL-6 (P=0.02) and leptin (P=0.002), pro-inflammatory mediators associated with cardiovascular risk. Levels of adiponectin, an adipokine with anti-inflammatory and anti-atherogenic functions, were significantly lower in the overweight group (P=0.002). The leptin/adiponectin ratio, a preferential atherogenic marker was significantly increased in women who are overweight (P=0.02). LBP, CRP, leptin, and adiponectin levels significantly correlated with BMI, but not with age and there was a significant correlation between LBP and IL-6 levels. These results reveal the presence of subclinical endotoxemia and a pro-inflammatory state in overweight women and are of interest for further studies with the goal for improved understanding of cardiovascular health risks in women.

## Introduction

Obesity and the closely related metabolic syndrome are associated with an increased risk for cardiovascular disease (CVD) and other debilitating and lethal disorders (1–6). Publicly available information on the World Health Organization (WHO) website states that in 2016 around 1.9 billion adults (people over 18 years of age) were overweight, and more than 600 million were obese and the expectations are that more than 2.16 billion people will be overweight and 1.12 billion will be obese by 2030. A major underlying factor driving the pathogenesis in obesity and metabolic syndrome is the presence of a chronic low-grade inflammation, which is characterized by increased circulating IL-6 and other cytokines, as well as altered levels of adipokines, such as leptin and adiponectin (3, 7–11). An important driver of the inflammatory state in obesity is *metabolic endotoxemia*, manifested by increased gut lipopolysaccharide (LPS)-containing microbiota and the consequent compromising of intestinal permeability that leads to increased circulatory LPS levels (12–15). Obesity-associated metabolic endotoxemia and chronic inflammation promote metabolic derangements and are associated with increased cardiovascular risk (9, 15–21).

In addition to obesity, there is evidence that overweight individuals may be at increased risk for CVD and other diseases (22, 23). CVD is the leading cause of death among women in the United States (6, 24). Increased cholesterol levels (hypercholesterolemia) and high blood pressure (hypertension) are important risk factors for CVD (6, 25). Elevated total cholesterol, hypertension, and excessive body weight have been linked to age-dependent increases of coronary heart disease incidence and mortality in both men and women, but to a larger extent in women (26). However, the underlying explanation for sex-specific differences in the CVD pathophysiology remain poorly understood (6). As recently summarized, “Cardiovascular disease in women remains understudied, under-recognized, underdiagnosed, and undertreated globally” (6).

While metabolic endotoxemia and inflammation have been documented in people with obesity and linked to CVD and other diseases, endotoxemia and inflammatory alterations in overweight individuals remain to be characterized. This is of specific interest for improved understanding of women’s cardiovascular health. To generate insight, we profiled a panel of plasma biomarkers of endotoxemia and inflammation, previously associated with CVD in obesity, in a cohort of women who were overweight compared to those who were lean. As dyslipidemia and hypertension are recognized leading traditional risk factors for CVD in women (6, 25), we enrolled subjects having high cholesterol and/or high blood pressure in both groups. We observed increased circulating levels of LBP, a marker of metabolic endotoxemia in parallel with elevated CRP, leptin, and IL-6 levels, and decreased adiponectin levels in overweight women.

## Materials and Methods

### Human subjects and samples

All methods were carried out in accordance with relevant guidelines and regulations. Frozen plasma was obtained from research subjects who participated in the Institutional Review Board (IRB)-approved Genotype and Phenotype (GaP) registry (http://www.gapregistry.org), a research program at the Feinstein Institutes for Medical research, Northwell Health. All research subjects completed an informed consent prior to study participation. The consent permits the use of specimens for future research. The study was approved by the Northwell Health IRB - IRB #09- 081A. Participants gave random blood samples and were chosen based on gender, BMI (lean: 18-24.9 kg/m^2^ (N=20) vs. overweight: 25-29.9 kg/m^2^ (N=20)), age, demographic information, and health/medical information (**Supplementary Table 1**). Subjects in both groups were relatively healthy, except they had self-reported hypertension (systolic ≥130mm Hg and diastolic >80 mm Hg) and/or high cholesterol (200 mg/dL) and minor conditions, including acne, eczema, gastroesophageal reflux disease (GERD), drug allergies, and other allergies, as well as osteoarthritis, osteopenia, osteoporosis. Excluded conditions were Lyme disease, cancer (solid and blood [leukemia, lymphoma, etc.]), anemia, pancreatitis, emphysema, asthma, chronic obstructive pulmonary disease (COPD), inflammatory bowel disease (ulcerative colitis, Crohn’s), lupus, rheumatoid arthritis, valvular disease, heart failure, HIV, excess alcohol use, diabetes (types 1 and 2) and Alzheimer’s disease and other neurological conditions that would impair the subjects’ ability to consent, as well as those using steroids, insulin, metformin, or glyburide and those who smoke or vape.

### Plasma sample analyses

All plasma samples were collected from consented GaP participants prior to the COVID-19 pandemic, aliquoted, and stored at -80°C in the Boas Center Biorepository. Just prior to analysis plasma samples were thawed and then assayed for numerous analytes (using dilutions optimized in prior studies) according to the manufacturer’s guidelines: adiponectin using the adiponectin/Acrp30 ELISA (DY1065, R&D System, lower limit of detection [LLoD] 15.6pg/ml), C-reactive protein or CRP by ELISA (DY1707, R&D Systems, LLoD 15.6pg/ml); leptin by ELISA (DY398-05, R&D Systems, LLoD 31.2pg/ml); LPS binding protein or LBP by ELISA (DY870-05, R&D Systems, LLoD 0.8ng/ml); and IL-6 using the V-plex MSD platform (K151QXD-2, Meso Scale Discovery, LLoD 0.06pg/ml).

### Statistical analysis

Data were analyzed using GraphPad Prism 9.5.1 software and applying an unpaired Student’s t test with Welch’s correction; P<0.05 was considered significant. The linear correlation for the data was analyzed using the Pearson’s correlation coefficient (or Pearson’s r). The strength of the correlations was assessed based on the r values and considered to be weak (0.2-0.39), moderate (0.40-0.59), or strong (0.6-0.79). Graphical representations were created using GraphPad Prism 9.5.1.

## Results

### Basic demographics of the study population

The average age and BMI for the lean and overweight cohorts is shown in **Table 1**. There was no statistically significant difference in subjects’ age. The average BMI of the subjects in the overweight group was significantly higher compared to the average BMI of the lean group (**Table 1**).

**Table 1.** Study participants’ age and BMI.

| Subjects | Lean | Overweight | p value |
| --- | --- | --- | --- |
| Age (years) | 55.65±9.1 | 59.7± 6.1 | P=0.12 |
| BMI (kg/m<sup>2</sup>) | 22.4±1.6 | 27.0±1.5 | P<0.0001 |

### Circulating markers of endotoxemia and inflammation are altered in overweight women

LBP is an important mediator of LPS interactions with immune cells and the LPS-induced transcription of pro-inflammatory cytokines (27). Because of the documented difficulties in measuring LPS in biological fluids (28), LBP has been proposed as a useful marker for activation of innate immune responses to microbial products, such as LPS (29–31). Increased levels of circulating LBP have been determined in obesity and the metabolic syndrome and associated with increased circulating IL-6 levels, impaired insulin resistance and cardiovascular risk (29, 31–33). In the present study we observed significantly increased plasma LBP in the overweight group compared with the lean group (**Figure 1A**). In addition, plasma levels of CRP, a general inflammatory marker, were significantly higher in the overweight women (**Figure 1B**), as were the cytokine IL-6 and the adipokine leptin (**Figure 1C, D).** In contrast, the levels of adiponectin were significantly lower in the overweight group **(Figure 1E).** In addition, the leptin/adiponectin ratio values were significantly increased in the overweight compared with the lean group (**Figure 1F).**

**Figure 1.**
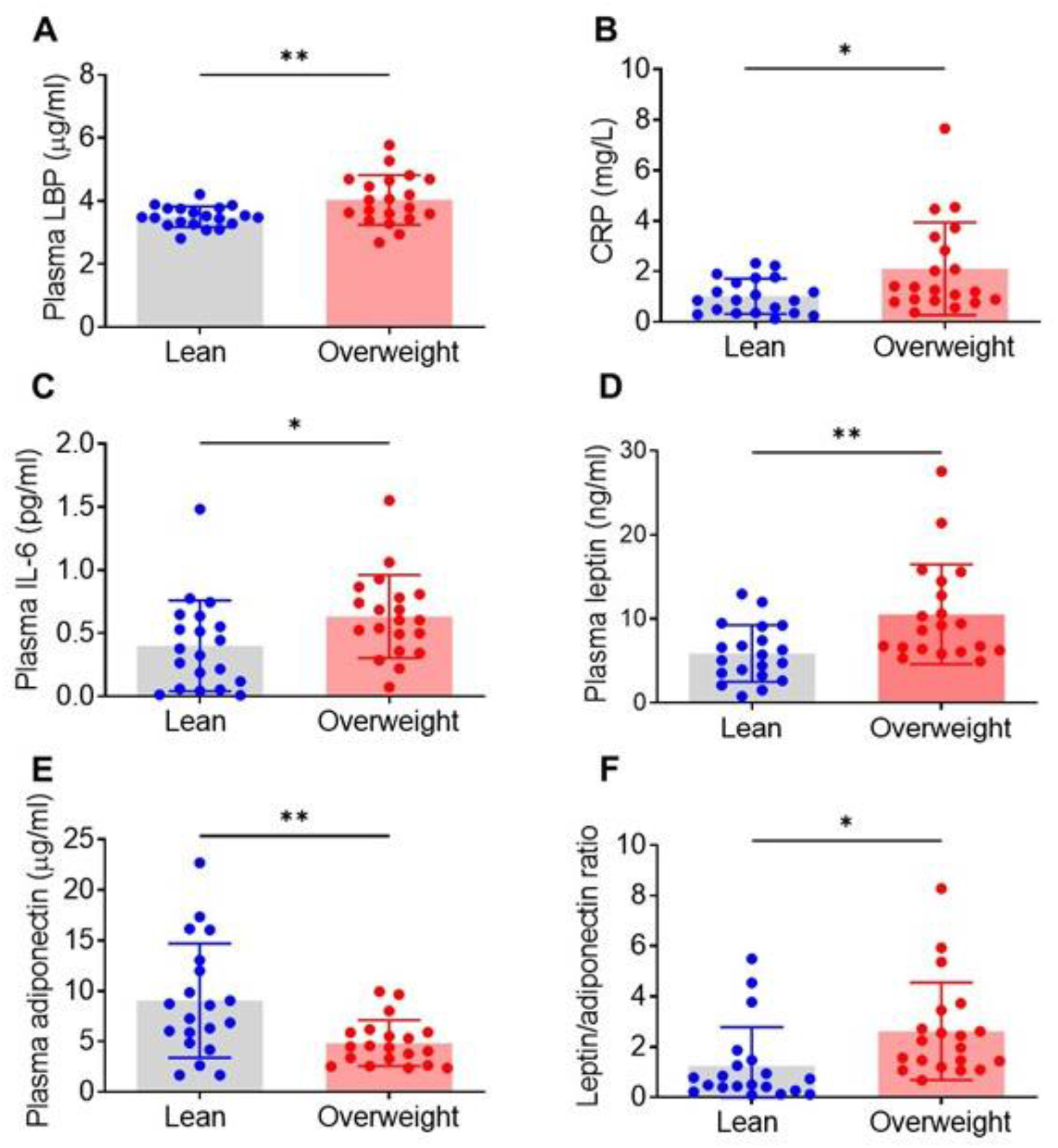
Levels of circulating markers of inflammation are altered in overweight women compared with lean women. Plasma samples of overweight and lean subjects were analyzed for (**A**) LPB, (**B**) CRP, (**C**) IL-6, (**D**) leptin, and (**E**) adiponectin as described in Materials and Methods, and leptin/adiponectin ratios (**F)** were calculated. Data are shown as mean ± SD (*P=0.01 (CRP; *P=0.02 (IL-6 and leptin/adiponectin ratio; **P=0.002 (leptin and adiponectin); **P=0.005 (LBP))

### Plasma markers of endotoxemia and inflammation correlate with BMI

Additional data evaluation revealed that plasma inflammatory marker alterations correlated with BMI of the study subjects (**Figure 2).** A moderate, but significant correlation was observed between plasma LBP and BMI (**Figure 2A**). Of note, it appeared that there was a breakpoint in the BMI vs. LBP relationship at the lower “overweight” limit. Similarly, we observed a moderate, but significant correlation between plasma CRP and BMI (**Figure 2B**). There was a weak, non-significant correlation between plasma IL-6 levels and BMI (**Figure 2C**). A moderate and significant correlation was observed between plasma leptin and BMI (**Figure 2D**) and between plasma adiponectin and BMI (**Figure 2E**).

**Figure 2.**
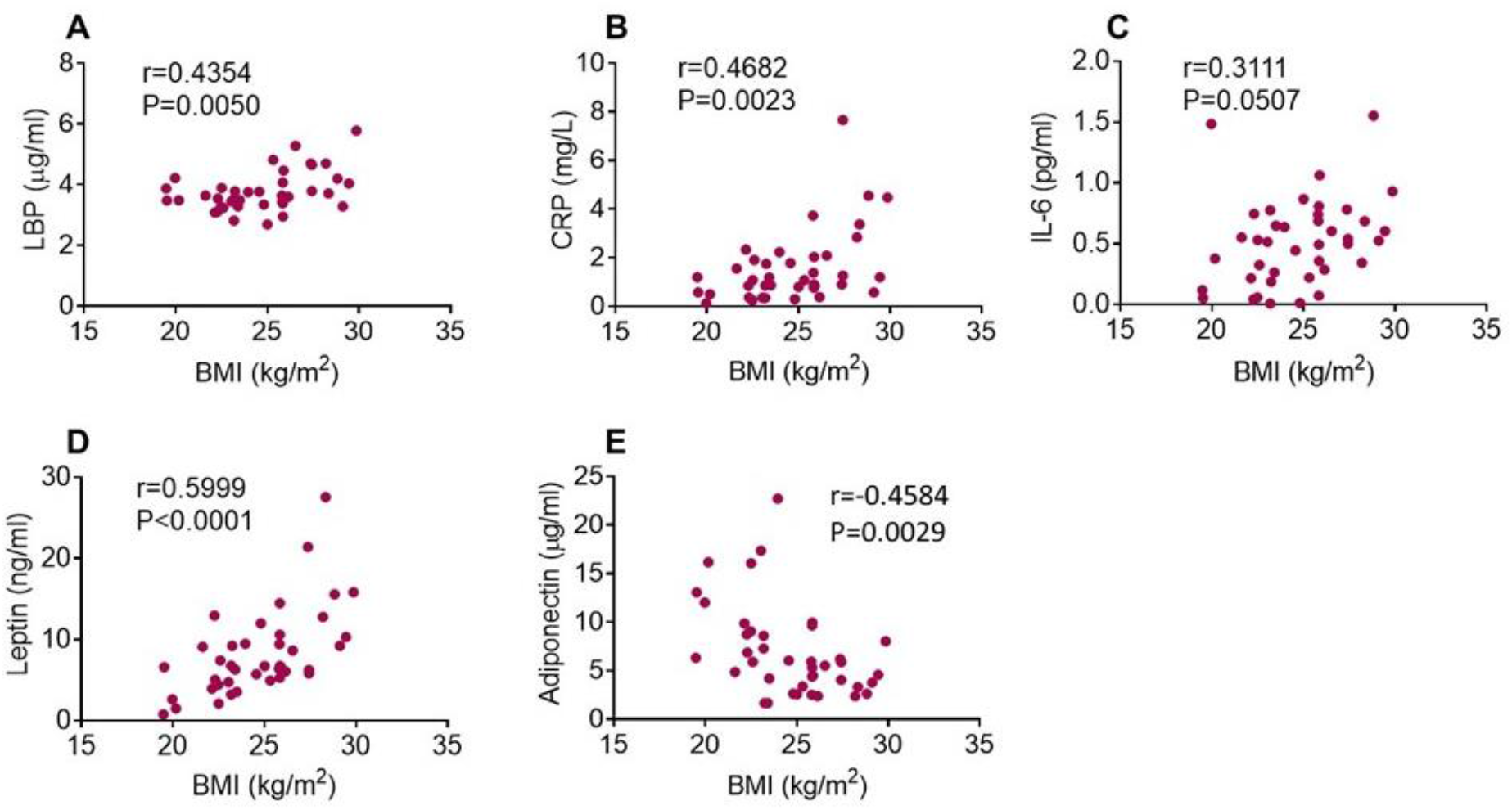
Correlation of plasma inflammatory indices and BMI. Plasma LBP (**A**), CRP (**B**), leptin (**D**), and adiponectin (**E**) levels significantly correlate with BMI as indicated by P values and Pearson correlation coefficients (r). Plasma IL-6 levels do not correlate with BMI (**C**).

### Plasma LBP level correlate with plasma IL-6 levels

Inter-correlation analysis of the analytes revealed that LBP was significantly correlated with IL-6 As shown in **Figure 3**. In addition, LBP, and CRP, as well as LBP and leptin correlational analyses generated P values of 0.053 and 0.085 respectively.

**Figure 3.**
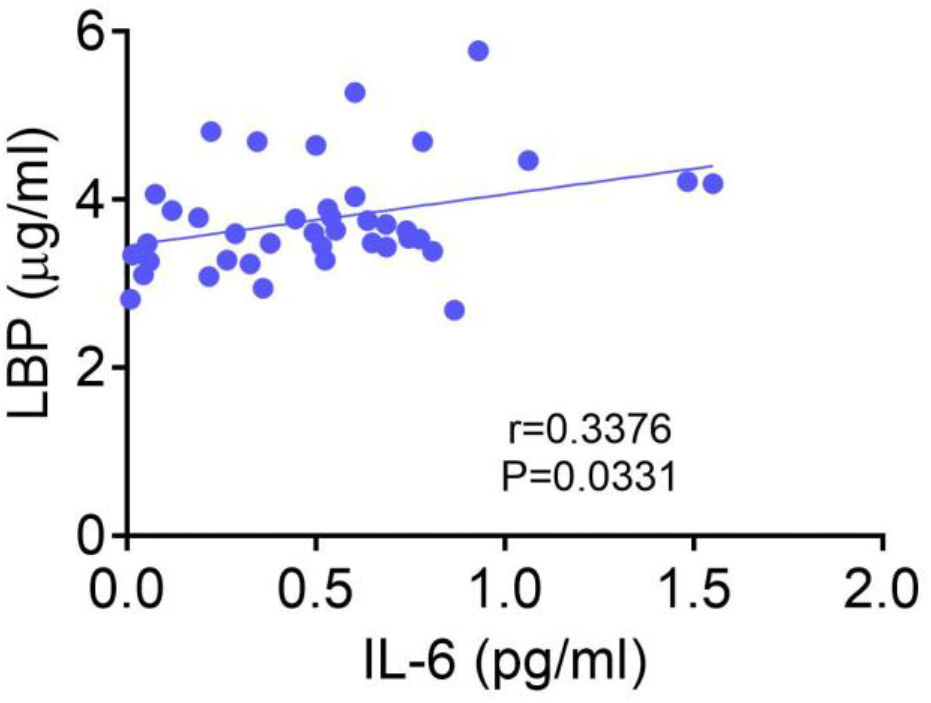
Correlation of LBP and IL-6. Plasma LBP levels significantly correlate with IL-6 levels as indicated by the P value and Pearson correlation coefficient.

Of note, no significant correlations were observed between the plasma inflammatory analytes and the age of the study participants as shown in **Supplementary Figure 1**.

## Discussion

Here, in women matched for age and having self-declared hypercholesterolemia and/or hypertension, we show that overweight individuals exhibited higher plasma concentrations of LBP and a pro-inflammatory state, indicated by increased levels of IL-6 and other proinflammatory markers and decreased adiponectin levels compared to lean controls. These differences are relatively small, but notable for the existence of a significant correlation with BMI as well as a positive correlation between LBP and the pro-inflammatory cytokine, IL-6.

Previously, the results of one large study evaluating 500 apparently healthy lean individuals, a majority being female, has shown that LBP is weakly, but highly significantly, correlated with BMI (34). In this study, LBP was also significantly positively correlated with blood pressure, HDL and LDL cholesterol, and triglycerides. Additionally, the results of a much larger (n=2,568 men and women) longitudinal study assessing the relationship between circulating LBP concentrations (the majority of which were within the normal range of less than 15 µg/mL) and cardiovascular risk factors show that the relationship between LBP and CRP, an accepted inflammatory marker, has a highly significant positive correlation (35). The current results conform to these findings and together indicate that markers of endotoxemia and a pro-inflammatory state vary continuously with the BMI, suggesting that adiposity of any degree, even if considered to be within the normal range, is associated with detectable levels of pro-inflammatory molecules. Whether these have clinical significance will require additional large, prospective clinical studies, but it stands to reason that a sustained low grade inflammatory milieu likely has clinical consequences.

Although there is a lack of data regarding the risk of subclinical endotoxemia and chronic inflammation in overweight individuals, previous studies in obese individuals have clearly indicated the presence of metabolic endotoxemia (based on LBP levels) and a chronic inflammatory state and their role in promoting further metabolic dysfunction and pathogenesis (3, 7–9, 36–42). Metabolic endotoxemia in obesity has been specifically linked to the pathogenesis of CVD (15, 21) and increased LBP levels have been directly associated with an increased risk of CVD (21). Endotoxemia increases the production of IL-6 and other cytokines and significantly contributes to a pro-inflammatory state. IL-6 and the adipokine leptin are key mediators of inflammation in obesity (3, 43). IL-6 has been characterized as an important link between obesity and coronary heart disease (44). Importantly, in a large prospective study, increased IL-6 levels were associated with a higher risk of CVD, specifically coronary heart disease, as strongly as major established risk factors, such as blood pressure and blood cholesterol levels (45) Of note, a significant correlation between plasma LBP and IL-6 levels has been previously documented in a large population spanning a large range of BMI and serum LBP concentrations (35), as well as subjects (males and females) with frank obesity (29). Leptin is an adipokine with an essential role in energy balance through a variety of functions, some of which are related to cardiovascular health (7, 18, 20). Increased leptin levels in obesity are associated with activation of pro-inflammatory signaling and increased thrombosis and arterial distensibility in obese patients (11, 46, 47). Elevated leptin levels arising from leptin resistance in obesity are associated with insulin resistance and CVD (18). In contrast, adiponectin is an adipokine with anti-inflammatory and antithrombotic properties (7, 48). Decreased plasma adiponectin levels were associated with an increased risk of myocardial infarction (49). Plasma levels of CRP (high-sensitive C-reactive protein), a general marker of chronic subclinical inflammation, have been positively correlated with plasma leptin levels and inversely with plasma adiponectin (50–52). The leptin/adiponectin ratio is indicated as a more reliable marker in CVD assessment compared with individual leptin and adiponectin measures (53–55) and proposed as a better marker of a first cardiovascular event in men than plasma leptin and adiponectin levels alone (56).

Until longitudinal data concerning subclinical endotoxemia are available, it may be prudent to institute proactive measures to monitor and reduce the circulating levels of LBP, as a surrogate biomarker for endotoxemia. As composition of the diet has been shown to be a critical driver of metabolic endotoxemia (reviewed in (57)), with high saturated fat ingestion causing postprandial endotoxemia with increases in IL-6 even in lean subjects (58, 59), dietary intervention would be a reasonable initial step. Other possibilities include pharmacological interventions, including the potential development of anti-LPS peptides which neutralize LPS signaling of immune system activation (57).

In addition to the small number of subjects assessed, there are several limitations of this study. An important one is that the degree of abdominal adiposity (in contrast to subcutaneous fat deposits) has been implicated in the pathogenesis of a systemic inflammatory response and correlates well with the production of pro-inflammatory cytokines (60). Of note, as a single measure BMI cannot fully capture this variable as it is quite insensitive to changes in regional body composition. That is, a smaller waist circumference for any given BMI is indicative of subcutaneous fat deposits, in contrast to an abdominal location in an individual possessing a larger waist circumference. A consensus has appeared that including the waist circumference as a measured variable provides information independent of the BMI, and when both are considered together the predictive accuracy of cardiometabolic risk is significantly increased (61). Additional limitations include other potentially contributing factors that were not assessed, including the existence of abnormal glucose tolerance, HOMA-IR information, lipid profile, degree of sedentary behavior, and the confirmation, severity and pharmacological treatment status of self-reported hypercholesterolemia and hypertension, among others. Lastly, as dietary factors are a major driver of the appearance of LPS into the circulation, in future studies plasma samples should be obtained under standardized fasting conditions.

## Conclusion

Our results indicate that despite the presence of hypertension and/or high cholesterol levels in two groups of women characterized using BMI as overweight versus lean, circulating biomarkers of innate immune activation and inflammation, including LBP, CRP, IL-6, and leptin are increased and adiponectin is decreased in the overweight group. Previous studies have focused on evaluating LBP and inflammatory markers in subjects with obesity and alterations of these molecules observed in obese individuals have been linked to an increased cardiometabolic risk.

Our study is among the first to provide insight into the overweight category and more specifically in overweight women. While the number of subjects in this study was small, the presence of subclinical endotoxemia and inflammation, the degree of which varied continuously with BMI encourage performing larger studies to better characterize individuals who are overweight, but not yet classified as obese. These individuals may benefit from therapy to alleviate chronic, low-grade inflammation as an additional risk factor for the development of CVD and other disorders.

## Acknowledgements

The authors wish to thank Gila Klein of the Genotype and Phenotype Registry, and the participants who provided blood for this study, as well as Michael Ryan and Bibu Jacob and staff members of the Boas Center Biorepository at the Feinstein Institutes for isolating and storing the plasma samples.

## Funding

This work was supported by the National Institutes of Health (NIH), National Institute of General Medical Sciences grants: RO1GM128008 and RO1GM121102 (to VAP) and R35GM118182 (to KJT).

## Data availability statement

All data generated or analyzed during this study are included in this published article and its supplementary information files.

## Declaration of interests

The authors declare that the research was conducted in the absence of any commercial or financial relationships that could be construed as a potential conflict of interest.

## Author contributions

CNM and VAP proposed the experimental concept and designed the experiments. XX and PKC performed the analyses. CNM, VAP, XX, PKC, and RPA analyzed data. CNM, VAP, and MB wrote and revised the manuscript. JR, PG, and KJT provided additional comments to finalize the paper.

---

**Supplementary Table 1.** Demographics of the study subjects.

| **Subject ID** | **BMI (kg/m<sup>2</sup>)** | **Age (years)** | **Ethnicity** | **Condition** |
| --- | --- | --- | --- | --- |
| **Lean Group** |  |  |  |  |
| 1 | 19.97 | 57 | Caucasian | HC* |
| 2 | 20.17 | 56 | Caucasian | Hypertension |
| 3 | 23.18 | 60 | Caucasian | HC |
| 4 | 22.31 | 56 | Caucasian | Hypertension, Other |
| 5 | 22.48 | 58 | Black/West Indian | HC |
| 6 | 19.53 | 63 | Caucasian | HC, Osteoporosis |
| 7 | 22.27 | 60 | Caucasian | HC, Other |
| 8 | 21.63 | 58 | Caucasian | HC |
| 9 | 22.50 | 61 | Caucasian | HC, Osteoarthritis |
| 10 | 23.49 | 62 | Caucasian | HC, Osteopenia |
| 11 | 23.96 | 63 | Black/African American | HC, Osteoarthritis |
| 12 | 24.56 | 47 | Caucasian | HC |
| 13 | 22.14 | 30 | Caucasian | HC |
| 14 | 23.18 | 46 | Caucasian/Latino/Hispanic | HC, Other |
| 15 | 22.59 | 58 | Caucasian | Hypertension, Other |
| 16 | 23.41 | 60 | Asian | HC, Hypertension, Other |
| 17 | 19.48 | 62 | Caucasian | HC |
| 18 | 23.24 | 61 | American Indian/Alaska Native | HC |
| 19 | 24.80 | 34 | Caucasian | HC |
| 20 | 23.04 | 61 | Caucasian | HC, Other |
| **Overweight group** |  |  |  |  |
| 21 | 28.34 | 68 | Caucasian | HC, Hypertension |
| 22 | 29.12 | 67 | Caucasian | HC, Hypertension |
| 23 | 28.83 | 53 | Black/West Indian | Hypertension |
| 24 | 25.84 | 52 | Caucasian | Hypertension |
| 25 | 26.54 | 66 | Caucasian | HC, Hypertension, Other |
| 26 | 29.86 | 56 | Caucasian | Hypertension |
| 27 | 25.32 | 57 | Caucasian | HC, Hypertension |
| 28 | 29.45 | 61 | Caucasian | HC, Hypertension |
| 29 | 25.82 | 66 | Caucasian | HC, Osteoporosis |
| 30 | 28.19 | 45 | Caucasian | HC, Other |
| 31 | 27.43 | 58 | Caucasian | HC |
| 32 | 25.88 | 66 | Caucasian | HC, Osteoarthritis |
| 33 | 25.84 | 68 | Caucasian | HC, Hypertension, Osteoarthritis, Other |
| 34 | 25.84 | 59 | Black/African American | HC |
| 35 | 26.15 | 64 | Caucasian | HC |
| 36 | 25.79 | 61 | Caucasian | Hypertension |
| 37 | 27.43 | 56 | Asian | HC |
| 38 | 25.01 | 53 | Caucasian | Hypertension |
| 39 | 25.85 | 61 | Caucasian | HC, Hypertension |
| 40 | 27.37 | 57 | Caucasian | HC, Hypertension |

\*HC (High cholesterol levels)

**Supplementary Figure 1.**
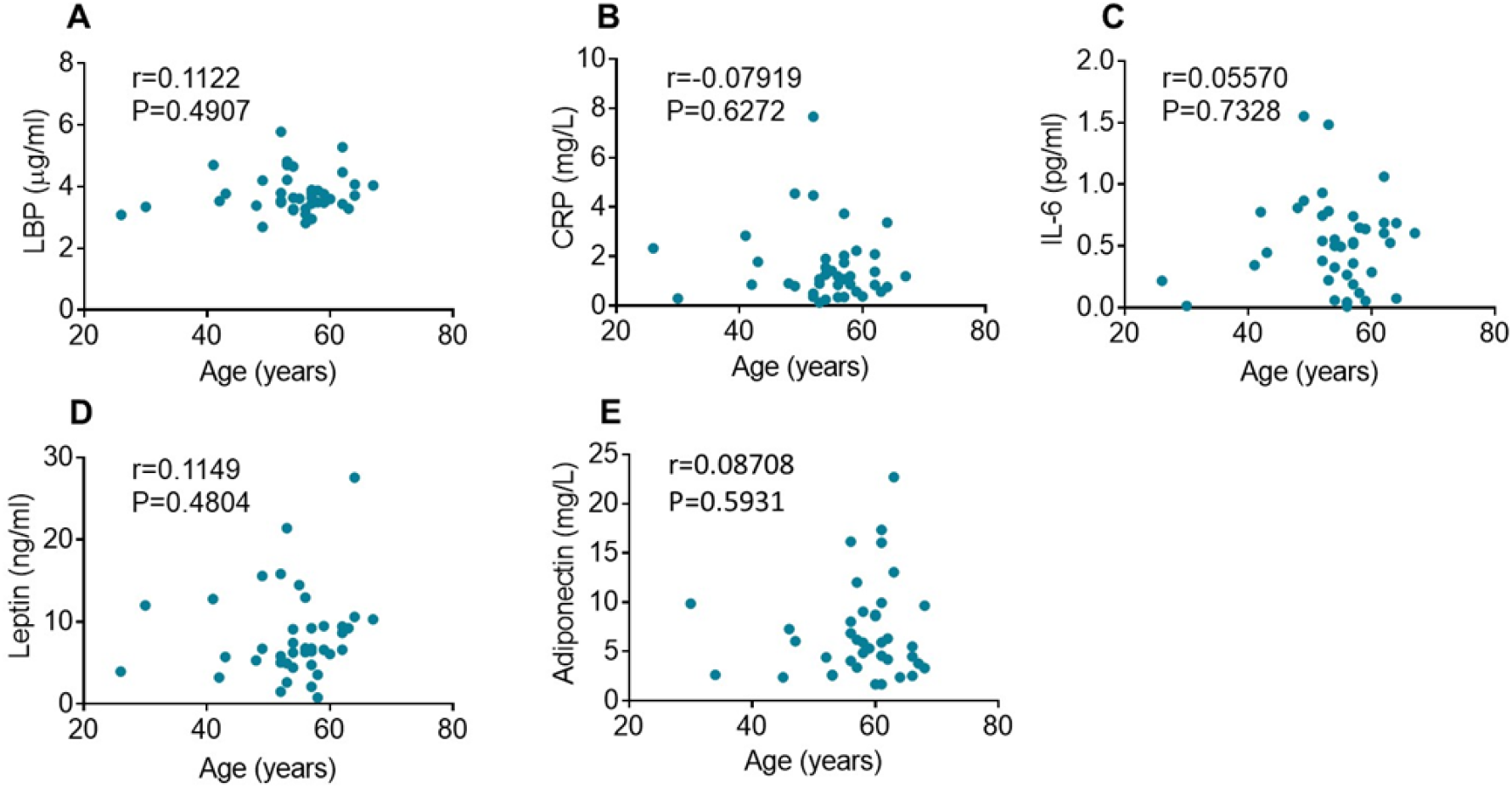
Correlation of plasma inflammatory indices and age. Plasma LBP (**A**), CRP (**B**), IL-6 (**C**), leptin (**D**), and adiponectin (**E**) levels do not significantly correlate with age as indicated by Pearson correlation coefficients (r) and p values.

## Notes

### Competing Interest Statement

The authors have declared no competing interest.

### Summary of Updates

Adding an author - Jesse Roth, MD; adding a figure - figure 3.

